# AstraPTM: Context-Aware PTM Prediction Model for Large-Scale Proteins

**DOI:** 10.1101/2025.02.20.639276

**Authors:** Aniruddh Goteti, Çağlar Bozkurt

## Abstract

Post-translational modifications (PTMs) are critical molecular events that profoundly influence protein stability, localization, and function. While numerous computational tools exist for PTM site prediction, most struggle with handling large proteins and rely on separate models for each modification. To address these challenges, we introduce **AstraPTM**, a transformer-based framework that predicts 25 PTMs in a single pass. By leveraging advanced protein embeddings (ESM2) and training on a high-coverage dbPTM dataset, AstraPTM captures both short-range sequence motifs and long-range interactions across proteins without a sequence length limitation.

AstraPTM combines a binary classification module—indicating whether a residue is modified—with a multi-label module that pinpoints specific PTM types. This dual approach achieves high accuracy on well-represented PTMs (e.g., phosphorylation, glycosylation) while maintaining sensitivity for rarer modifications. In benchmarks against existing methods such as MusiteDeep and MIND-S, AstraPTM demonstrates competitive or superior performance, demonstrating AUC-ROC above 99% for well-represented modifications, underscoring its versatility for proteome-wide annotation. Beyond prediction, the model’s capacity to handle full-length proteins offers a powerful resource for researchers investigating PTM crosstalk and disease pathways, ultimately bridging the gap between large-scale omics data and targeted biomedical applications.

## Introduction

Post-translational modifications (PTMs) are chemical changes occurring after protein synthesis, significantly influencing a protein’s stability, localization, and interactions (1; 2; 3). Common examples include phosphorylation (adding a phosphate group), glycosylation (adding sugar molecules), and ubiquitination (tagging proteins for degradation). By altering how proteins fold and function, PTMs modulate nearly every aspect of cellular biology and can play pivotal roles in disease progression (4; 15).

Researchers have developed numerous computational tools to predict PTM sites from protein sequence and structural cues. Some approaches focus on proximity-based features, while others draw on evolutionary history to highlight residues likely to undergo modification (6; 7; 2). Understanding these prediction methods is especially relevant in medical contexts, where aberrant PTM patterns are frequently implicated in cancers, neurodegenerative disorders, and other pathologies. By pinpointing these abnormal sites, scientists can design targeted therapies aimed at restoring or inhibiting specific modifications (8; 9; 10).

Despite notable progress, many existing tools face limitations such as handling large proteins, accommodating highly imbalanced PTM data, or requiring separate models for different modification types. To address these gaps, we introduce **AstraPTM**: a context-aware, transformer-based model capable of predicting 25 PTMs in a single pass. By combining comprehensive protein embeddings, sequence-level context, and extensive training data, AstraPTM aspires to deliver more robust and scalable PTM predictions for both research and clinical applications.

## 2 Literature Review

Post-translational modifications (PTMs) are key biochemical processes that modify proteins after synthesis, influencing their stability, localization, and interactions (11; 15; 2). These modifications include phosphorylation, glycosylation, ubiquitination, and acetylation, each capable of reshaping a protein’s structure and function. Given their dynamic and context-dependent nature, predicting PTMs is vital for understanding protein regulation and complex cellular pathways.

Early computational tools such as **SAPH-ire** leveraged structural data (e.g., solvent accessibility) and family-level modification counts to predict PTM sites (11). Subsequent models like **Sitetack** demonstrated that incorporating known PTM locations and co-occurrence patterns (e.g., O-glycosylation with phosphorylation) can further enhance predictive accuracy (15). These advances underscore the importance of analyzing PTM networks holistically, as co-evolving PTMs may have synergistic effects on protein function (2).

Research has also highlighted the clinical significance of PTM prediction, particularly in cancer and neurodegenerative diseases, where abnormal PTM profiles can reveal therapeutic targets (1; 16). By identifying key phosphorylation or ubiquitination sites, for instance, drug design can focus on inhibiting or modifying these PTMs, thereby improving treatment specificity and efficacy. Tools like **jEcho** and **PEIMAN** have further integrated sequence, structure, and evolutionary data to generate user-friendly platforms for PTM visualization and enrichment analysis, assisting researchers in systematically assessing PTM implications (17; 10).

Recent developments in **machine learning** and **AlphaFold**-driven structural predictions have greatly enhanced PTM site prediction (18). By combining high-resolution structural data with multi-scale sequence representations, these models offer more nuanced insights into PTM location and function. This synergistic approach signifies a leap forward in our capacity to decode complex regulatory networks governed by PTMs, thereby paving the way for improved therapeutic interventions and a deeper comprehension of protein biology.

Recent PTM prediction models have demonstrated notable improvements over established methods (12; 13; 14; 15), yet they still exhibit considerable shortcomings. One key issue is the reliance on limited context windows, which restricts the ability to capture long-range dependencies crucial for context-sensitive post-translational modifications. Additionally, many models do not include a sufficiently large or diverse set of proteins in their training corpora, hampering the identification of nuanced PTM occurrence patterns across different organisms and protein families. Finally, most current approaches lack an open-set or generative strategy, focusing only on PTMs already established by experimental evidence and rarely venturing beyond them to predict entirely novel modifications.

In summary, PTM prediction research continues to expand, encompassing structural, sequence-based, and evolutionary strategies. The integration of these data streams, coupled with advances in deep learning, positions PTM analysis as an essential component of modern proteomic research and drug discovery efforts.

## 3 Keys to Successful PTM Prediction

**Careful dataset design** is critical for achieving successful predictions with deep learning. When predicting **post-translational modifications (PTMs)**, the following considerations are especially important:

### Contextual Awareness

PTM occurrence heavily depends on the surrounding protein sequence, so incorporating sufficient context around each residue is crucial for accurate prediction.

### Feature Enrichment

To help the model learn the nuances of PTM sites, additional biochemical and structural properties should be included in the feature set. This enrichment goes beyond raw sequence data to capture meaningful context.

### Dataset Size and Diversity

While a dataset should be homogeneous in format, it must also represent the various

PTMs found in real-world data. Ensuring *both* sufficient size and diversity allows the model to generalize effectively to different PTMs.

## 4 AstraPTM: Steps to a Context-Aware Model

AstraPTM is a deep learning framework designed to predict which residues carry post-translational modifications (PTMs) *and* to specify the type of PTM at each site. By leveraging powerful **ESM2 embeddings** and a **Transformer** architecture, AstraPTM captures both local and global sequence information without being constrained by fixed windows. This enables accurate predictions for proteins of varying lengths—without a limitation in length—while efficiently learning from multiple types of PTM data.

### Highlights of the AstraPTM Approach

#### Comprehensive Context Capture

Unlike models that rely on fixed-size windows, AstraPTM’s transformer-based design uses self-attention to integrate information across the full protein. Coupled with ESM2 embeddings, this allows the model to detect subtle, long-range dependencies crucial for accurate PTM predictions.

#### Scalability for Large Proteins

By avoiding manual sequence truncation, AstraPTM can handle proteins without a limit in sequence length. Its attention mechanism dynamically scales to capture interactions among distant regions.

#### Binary & Multi-label Synergy

A single framework handles both the identification of PTM sites (*binary task*) and the classification of specific PTM types (*multi-label task*). This approach improves overall robustness, especially for rare or novel modifications.

### Benefits of Having Binary and Multi-label Models

For most PTMs, the scarcity of experimental data constrains the predictive performance of machine learning methods. As a result, many machine learning models designed for PTM prediction restrict the range of PTMs they support within their multi-label frameworks.

To overcome this limitation and assist researchers in identifying novel or rare PTMs in their proteins of interest, we frame the PTM prediction task around two closely related questions:

1. Where do PTMs occur along the protein sequence?
2. Which types of PTMs are likely to be present at those locations?

This approach stems from the limited availability of PTM data. The dbPTM database, for example, shows a highly imbalanced distribution: only 25 PTMs appear in more than 100 proteins, and just 14 exceed 1,000 annotated proteins. Consequently, a single large multi-class model spanning all 72 dbPTM PTMs would struggle to learn meaningful distinctions among underrepresented PTM types.

By contrast, a **binary classification model** can effectively learn to detect whether a residue is modified at all—even if a specific PTM is rare or unobserved. The companion **multi-label classification model** then refines the prediction by specifying which PTM(s) are most likely to be present at that site. In practice:

#### Binary PTM Presence

The model can potentially capture **novel** or **rare** PTMs by leveraging shared contextual cues across different PTMs.

#### Multi-Label PTM Type

Where sufficient data exists, the model can discriminate among known PTM categories for more detailed predictions.

Together, these models enhance both accuracy and robustness, balancing broad detection with targeted classification.

### Contextual Awareness via ESM2 and Transformers

To capture the rich context crucial for PTM prediction, **four key steps** define AstraPTM’s pipeline:

#### 1. Incorporated ESM2 Embeddings

Each residue is encoded using the `esm2_t33_650M_UR50D` variant, which provides a 1,280-dimensional embedding trained on large-scale protein data. These embeddings contain evolutionary and structural signals that give AstraPTM a strong starting point for PTM discrimination.

#### 2. Transformer Architecture with Multiple Attention Heads

Building on ESM2’s rich embeddings, we use a transformer with **eight attention heads** per layer. Multiple heads let the model focus on distinct “subspaces” of protein features, from local sequence motifs to long-range structural interactions. This goes beyond raw embeddings by learning how specific residues and neighboring regions influence PTM sites.

#### 3. Full-Length Sequences

Rather than trimming proteins, we train on entire sequences (up to 3,000+ residues). This ensures that any remote interactions—which may be critical for PTM formation—are retained. Such a full-sequence approach outperforms methods restricted to short windows.

#### 4. Contextual Labeling with PTM Data

We curated a labeled dataset from dbPTM, annotating each residue with the PTM type and relevant structural context (when available). These detailed labels teach the transformer to distinguish which environmental cues correlate with specific modifications.

**Figure 1:**
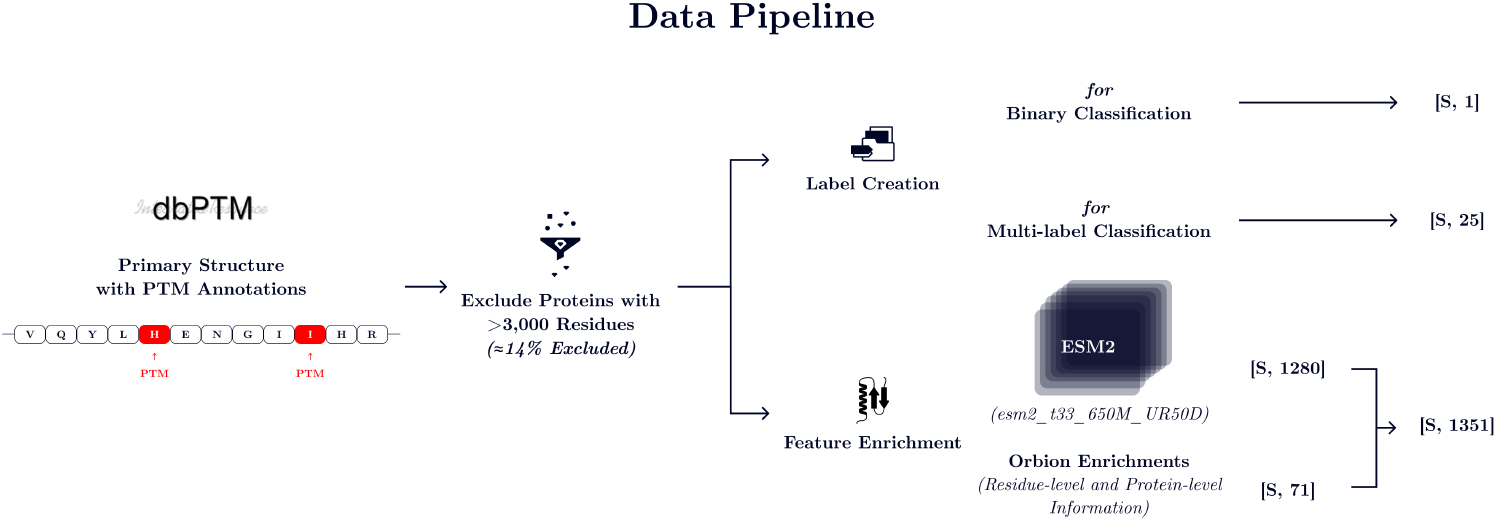
A diagram visualizing how the features and the labels of AstraPTM model arestructured.

### Training Data Structure

We leveraged **dbPTM** as our primary data source, which initially contains over **280**,**000 unique proteins**. To manage computational constraints, we excluded proteins exceeding **3**,**000 residues**, removing approximately 40,000 entries. This filtering struck a balance between covering a wide range of proteins and maintaining tractable sequence lengths for model training.

For each protein in the filtered dataset, we generated **ESM2 embeddings** (see ‘Contextual Awareness”) to capture evolutionary and structural signals. To further enrich these representations, we incorporated additional **residue-level** and **protein-level** features that together form a **1**,**351-dimensional** input per residue:

#### Residue-level Information

Hydrophobicity, Bulkiness, Karplus-Schulz Flexibility Index, Atomical Composition, Side Chain Polarity Index, Charge at pH 7.0, Aromaticity, Van der Waals Radii, Number of Total Bonds

#### Protein-level Information

Sequence Length, Molecular Weight, Instability Index, Gravy Score, Secondary Structure (% *α*-Helices, % *β*-Sheets, % Coils), Molar Extinction Coefficient, Isoelectric Point, Charge at Different pH Points, Atomical Composition

By combining the **contextual richness of ESM2** with these specialized physicochemical and structural features, the final dataset ensures that our transformer-based model receives both **local** (residue-specific) and **global** (protein-level) contextual information. We divided this enriched dataset into **training, validation, and test splits** to reliably measure performance and guard against overfitting, thereby supporting robust PTM prediction and generalization to novel proteins.

### Data Details

#### Binary Classification

We included the data from dbPTM for **72 PTMs** in a broad “PTM vs. no PTM” classification. Even underrepresented modifications can inform the binary model, enhancing sensitivity to rare but potentially important PTMs. The PTMs included in the model (but not limited to): Phosphorylation, Ubiquitination, Acetylation, N-linked Glycosylation, Succinylation, Methylation, O-linked Glycosylation, Malonylation, Sulfoxidation, S-palmitoylation, Sumoylation, S-nitrosylation, Glutathionylation, Amidation, Hydroxylation, Neddylation, Pyrrolidone-carboxylic-acid, Glutarylation, Oxidation, Gamma-carboxyglutamic-acid, Lactylation, Crotonylation, Myristoylation, Sulfation, Formylation, C-linked Glycosylation, and others.

#### Multi-label Classification

For the multi-label classification task, we restricted our dataset to PTMs with a higher number of annotated protein observations. In particular, we included only those PTMs with more than 100 protein annotations; of these, 14 had more than 1,000 protein observations. Consequently, the multi-label model training set comprised PTMs such as Phosphorylation, Ubiquitination, Acetylation, N-linked Glycosylation, Methylation, Succinylation, Malonylation, O-linked Glycosylation, S-palmitoylation, Sulfoxidation, and several others.

**Table 1:** Summary of the total number of proteins that are used in training, test, and validation, in each category for both binary and multi-label classifications, separated by number of residues.

| Proteins | Binary Classification | Multi-label Classification |
| --- | --- | --- |
| Total | 95,789 | 95,417 |
| <100 Residues | 5,762 | 5,722 |
| 100-500 Residues | 52,561 | 52,319 |
| 501-1,000 Residues | 26,922 | 26,852 |
| >1,000 Residues | 10,544 | 10,524 |

## 5 Model Architecture

**AstraPTM’s architecture** is a sequence-to-sequence Transformer that takes, as input, a batch of protein residues—each residue described by **1**,**351 features**—and outputs logits indicating the likelihood of a PTM (for binary classification) or the likelihood of each possible PTM type (for multi-label classification).

**Figure 2:**
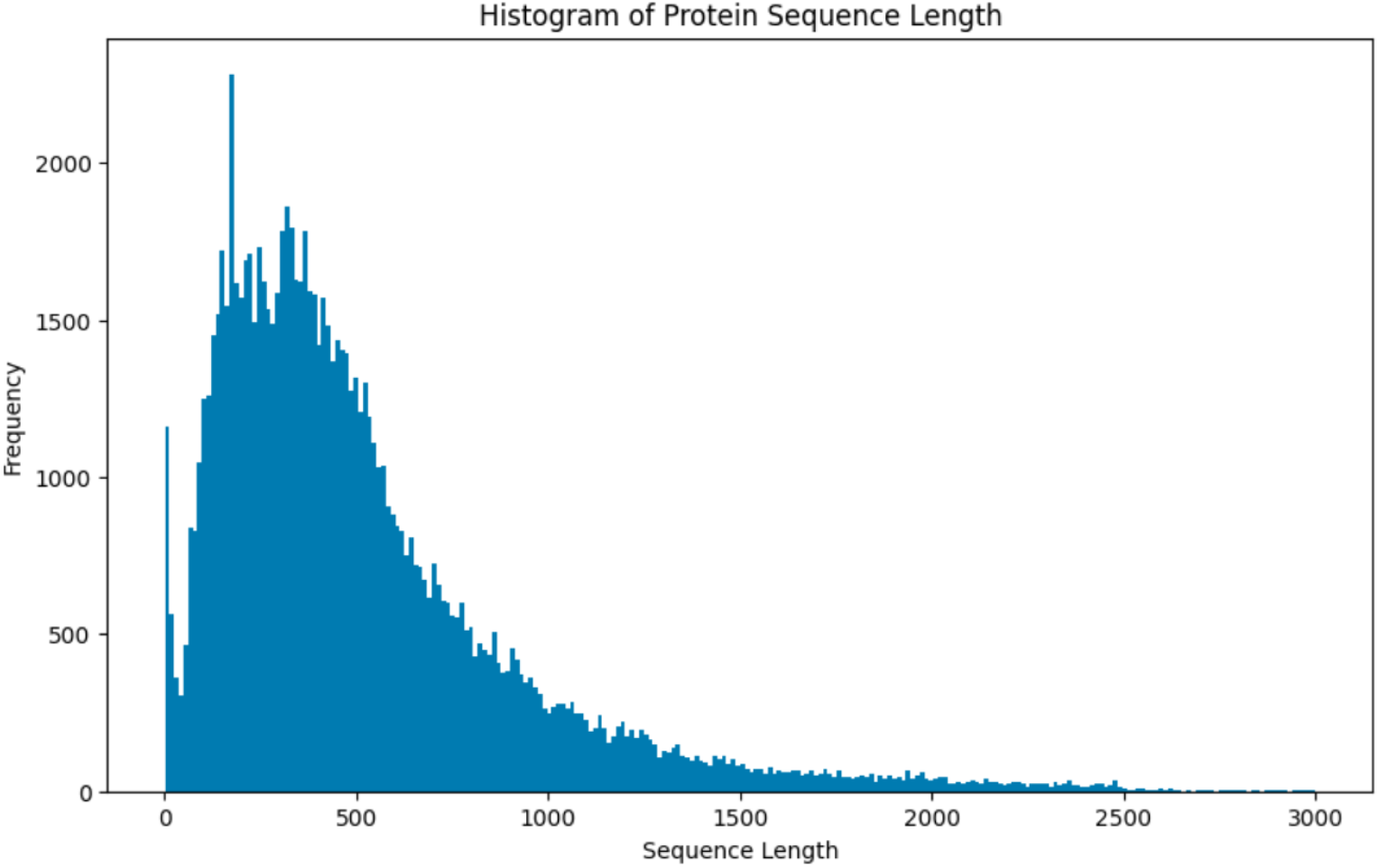
The histogram of the sequence length of the proteins that are included in the training of the model.

### Input Projection

A linear layer projects the 1,351-dimensional input to a hidden dimension of 512, compressing raw features into a space compatible with the transformer encoder.

### Positional Encoding

We add sine-cosine positional embeddings to each residue’s input vector, helping the model track ordering across the protein sequence.

### Transformer Encoder

We stack four layers, each with:

- 8 self-attention heads
- A feedforward network (4 × hidden dim)
- Layer normalization and dropout

This encoder can capture both short- and long-range dependencies critical for PTM prediction.

### Final Linear Layer

Maps the final hidden dimension (512) back to:

- 1 output for binary classification (PTM vs. no PTM),
- 25 outputs for multi-label classification (which PTM(s) at each residue).

**Figure 3:**
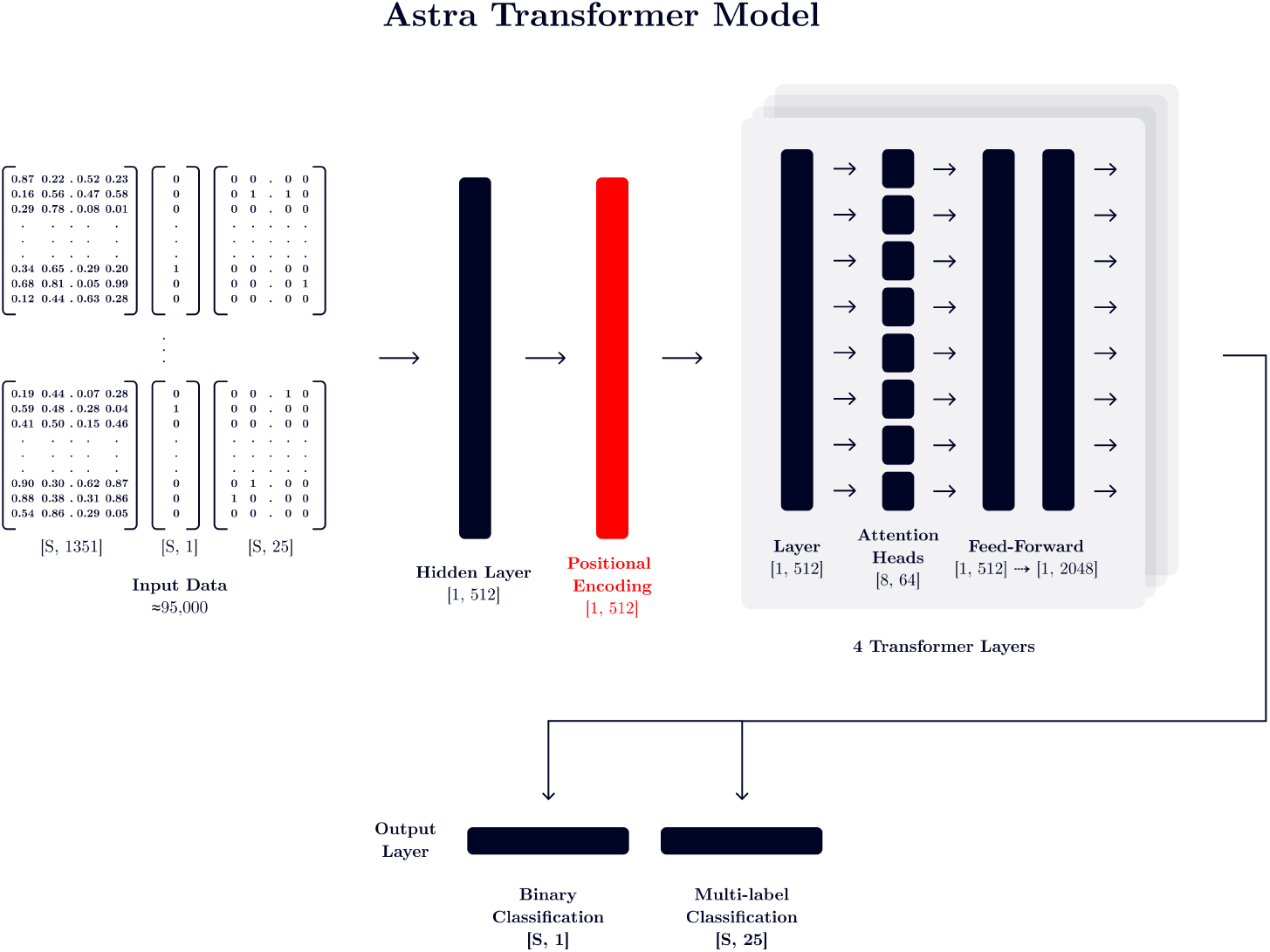
Diagram of the AstraPTM Transformer architecture for PTM prediction.

## 6 Results

AstraPTM’s performance was evaluated in two key modes: **binary** classification (whether a residue is modified at all) and **multi-label** classification (which PTM types are present). These complementary approaches are especially relevant for structural biologists and bioinformaticians who often need both a high-level scan for potential PTM sites *and* a fine-grained view of specific modifications that might alter protein conformation or function.

### 6.1 Overall Performance

We assessed AstraPTM using common metrics such as **AUC-ROC, AUC-PR, MCC**, Precision, Recall, F1 score, and Accuracy. Table 2 summarizes the high-level results for each classification task.

**Table 2:** High-level performance metrics for AstraPTM in binary and multi-label (25 PTMs) modes.

| Metric | Binary Classification | Multi-label (25 Labels) |
| --- | --- | --- |
| AUC-ROC | 96.18% | 99.57% (Micro) |
| AUC-PR | 94.46% | 85.29% (Micro) |
| MCC | 78.65% | 37.78% (Macro) |
| Precision | 86.33% | 72% (Weighted), 68% (Micro), 46% (Macro) |
| Recall | 89.71% | 84% (Weighted), 84% (Micro), 56% (Macro) |
| F1 | 87.98% | 77% (Weighted), 75% (Micro), 49% (Macro) |
| Accuracy | 89.46% | 94.3% (Weighted), 98.99% (Micro), 98.99% (Macro) |

#### 6.1.1 Binary Classification

This model exhibits high accuracy and a strong balance between precision (≈ 86%) and recall (≈ 90%). It effectively identifies whether a residue is modified or not, providing reliable guidance for downstream experiments or protein structure determination. By correctly flagging the majority of modified residues, the binary model offers a reliable “first pass” screen, minimizing missed sites while maintaining good specificity.

#### 6.1.2 Multi-label Classification

The multi-label model demonstrates excellent performance overall when aggregating all PTM labels (e.g., Micro AUC-ROC of ≈ 99.57%). However, lower macro metrics (MCC of ≈38%, F1 of ≈49%) reflect variability across individual PTMs, particularly those with fewer training examples.

Table 3 details the per-PTM results, highlighting that labels with sufficient data (e.g., Phosphorylation, N-linked Glycosylation) achieve strong metrics, whereas rare PTMs (e.g., Formylation, Neddylation, Lactylation) show high AUC-ROC but very low AUC-PR and MCC. These discrepancies underscore the impact of class imbalance in multi-label prediction.

**Table 3:** Per-PTM performance metrics for the multi-label model.

| Label | AUC-ROC | AUC-PR | MCC | Precision | Recall | F1-score | Accuracy |
| --- | --- | --- | --- | --- | --- | --- | --- |
| Formylation | 99.00% | 3.75% | 0.00% | 7.00% | 8.00% | 7.00% | 99.98% |
| Phosphorylation | 97.95% | 93.67% | 85.46% | 85.00% | 94.00% | 90.00% | 93.92% |
| Pyrrolidone-carboxylic-acid | 99.99% | 90.56% | 81.35% | 93.00% | 76.00% | 84.00% | 99.99% |
| Neddylation | 92.53% | 2.78% | 0.00% | 6.00% | 5.00% | 5.00% | 99.90% |
| Gamma-carboxyglutamic-acid | 99.99% | 70.72% | 43.17% | 65.00% | 73.00% | 69.00% | 99.99% |
| Sulfation | 99.34% | 35.14% | 9.90% | 46.00% | 61.00% | 53.00% | 99.99% |
| Hydroxylation | 99.40% | 61.25% | 62.52% | 68.00% | 61.00% | 64.00% | 99.96% |
| Oxidation | 99.53% | 3.18% | 0.00% | 10.00% | 8.00% | 9.00% | 99.98% |
| Succinylation | 98.65% | 30.74% | 9.20% | 34.00% | 44.00% | 38.00% | 99.02% |
| Amidation | 99.90% | 97.26% | 93.10% | 92.00% | 95.00% | 93.00% | 99.98% |
| Methylation | 97.72% | 46.50% | 41.83% | 54.00% | 41.00% | 47.00% | 99.49% |
| Malonylation | 97.64% | 16.10% | 0.00% | 19.00% | 35.00% | 24.00% | 98.97% |
| Glutathionylation | 99.46% | 35.95% | 10.13% | 37.00% | 53.00% | 44.00% | 99.78% |
| O-linked-Glycosylation | 94.40% | 24.67% | 17.94% | 32.00% | 36.00% | 34.00% | 99.30% |
| Sumoylation | 98.34% | 26.42% | 25.87% | 29.00% | 32.00% | 31.00% | 99.71% |
| S-palmitoylation | 99.46% | 59.56% | 53.31% | 58.00% | 49.00% | 53.00% | 99.80% |
| Sulfoxidation | 99.42% | 55.87% | 39.93% | 50.00% | 62.00% | 55.00% | 99.73% |
| N-linked-Glycosylation | 99.68% | 96.84% | 94.75% | 97.00% | 93.00% | 95.00% | 99.91% |
| GPI-anchor | 100.00% | 83.89% | 40.82% | 92.00% | 67.00% | 77.00% | 100.00% |
| Glutarylation | 99.18% | 14.91% | 0.00% | 26.00% | 15.00% | 19.00% | 99.97% |
| S-nitrosylation | 99.65% | 38.61% | 16.89% | 37.00% | 56.00% | 44.00% | 99.78% |
| Myristoylation | 99.92% | 87.46% | 85.35% | 93.00% | 85.00% | 89.00% | 100.00% |
| Acetylation | 97.59% | 61.20% | 48.78% | 56.00% | 68.00% | 62.00% | 96.18% |
| Lactylation | 98.66% | 0.85% | 0.00% | 1.00% | 22.00% | 2.00% | 99.55% |
| Ubiquitination | 94.27% | 64.80% | 54.30% | 52.00% | 70.00% | 60.00% | 91.90% |

### Key Insights

#### High-Confidence PTMs

Phosphorylation, N-linked Glycosylation, Myristoylation, and several others exhibit strong metrics (AUC-PR, MCC), making them prime targets for experimental validation (e.g., mutagenesis, structural analysis).

#### Moderate-Confidence PTMs

Acetylation or Ubiquitination can be influenced by data imbalance in certain families; further structure-based checks (e.g., solvent exposure) can refine predictions.

#### Rare/Underrepresented PTMs

Formylation, Neddylation, and Lactylation suffer from low Precision/Recall or near-zero MCC, indicating limited positive examples. Users should treat these predictions with caution and prioritize experimental followup.

Overall, the binary model reliably identifies PTM sites, while the multi-label model excels in detecting well-represented PTMs and provides potential leads for rare modifications. In practice, combining both models helps guide scientists on which sites to investigate further.

### 6.2 Practical Takeaways

#### Binary + Multi-label Synergy

The two-step approach helps *structural labs* focus on residues for biochemical assays and *bioinformatics* teams plan more targeted in silico analyses. Residues flagged as “modified” in the binary model but missing strong multi-label signals could represent novel or rare modifications.

#### Downstream Experimental Value

High recall means labs are less likely to miss crucial modification sites, crucial for large or multi-domain proteins.

#### Focus on Strong PTMs

If certain PTMs (phosphorylation, glycosylation) are especially relevant, AstraPTM’s high precision can accelerate site-directed mutagenesis or cryo-EM studies.

#### Guiding Structure Validation

Overlaying AstraPTM predictions on 3D models (e.g., AlphaFold, X-ray structures) can reveal if predicted sites are solvent-exposed or near active sites, informing functional hypotheses.

#### Rare PTMs

For modifications with sparse data, users may adopt additional validation strategies (mass spectrometry, literature cross-checks) to confirm or refute predicted sites.

## 7 Comparisons

As there are no other binary PTM prediction models aiming to answer the generic question, “where do PTMs occur along the sequence, regardless of PTM type?”, we focus on **multi-label** comparisons with popular methods.

AstraPTM addresses multiple PTMs simultaneously, whereas most other predictors traditionally focus on binary classification (one PTM at a time). We compare **AstraPTM** to **MusiteDeep, ModPred, Sitetack, MIND-S**, and a fine-tuned GPT-2 model (**PTMGPT2**), covering both classical and recent deep-learning approaches.

### Scope of Comparison

#### Sitetack, ModPred

Separate binary classifiers per PTM, potentially more finetuned but burdensome to deploy collectively.

#### MusiteDeep

CNN-based deep learning for specific PTMs, uses sliding windows.

#### MIND-S

Incorporates structural information (AlphaFold/PDB) for multi-label PTM analysis, but can be slow or require ensembles.

#### PTMGPT2

Uses a GPT-2 language model for multi-label PTM prediction but can be limited by sequence length constraints (~1024 tokens).

### Dataset and Model Nuances

While the comparisons showcase how models perform against each other, there are certain points that needs to be considered for a fair evaluation:

- The models **Sitetack** and **ModPred** are a set of binary classification models, meaning for every PTM, there is another model that needs to be deployed.
- Not all metrics of all models are publicly available, hence, although some comparisons are possible, the comparisons don’t reflect the full picture.
- **Not all models are trained and evaluated with the same dataset**. Hence, the comparisons are not completely reflective.
- The size of the dataset or the limitations on the dataset, such as protein length, may lead to misleading results, as it may limit models’ performances to only certain types of proteins.

Table 4 overviews each model’s dataset size, sequence length limits, PTM coverage, and classification type. Note that AstraPTM’s ability to handle longer proteins and unify multiple PTMs in a single model distinguishes it from older, window-based tools.

**Table 4:**
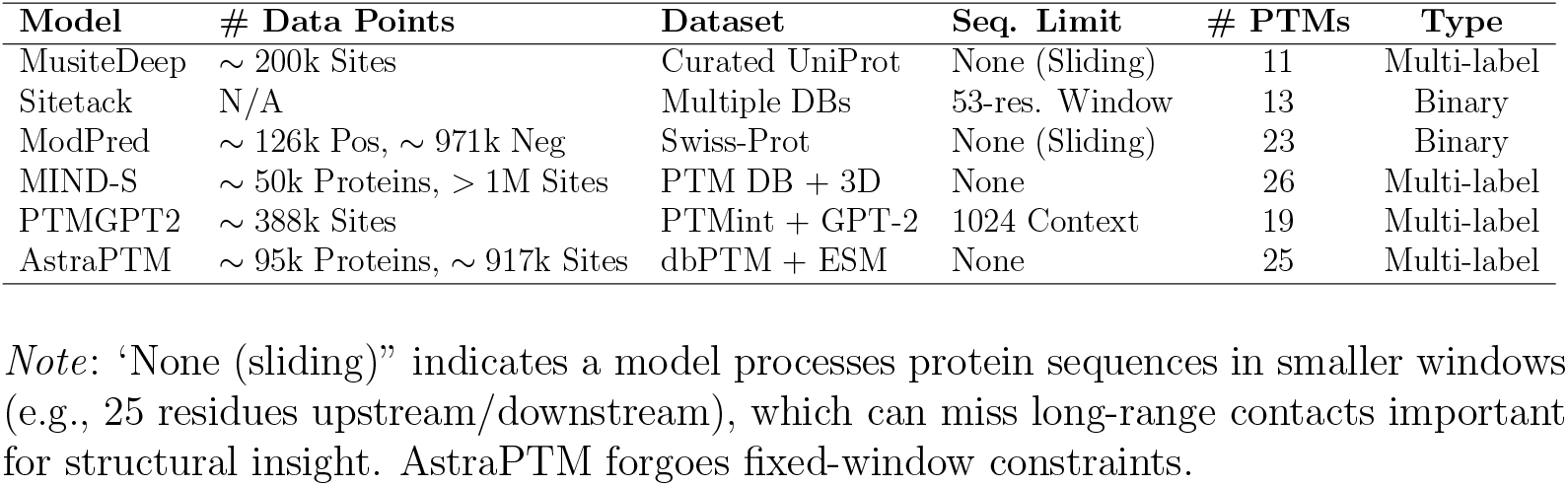
Comparison of PTM Prediction Models.

*Note*: ‘None (sliding)” indicates a model processes protein sequences in smaller windows (e.g., 25 residues upstream/downstream), which can miss long-range contacts important for structural insight. AstraPTM forgoes fixed-window constraints.

### 7.1 PTM-Specific AUC-ROC and AUC-PR

Tables 5 and 6 compare AstraPTM to MusiteDeep, ModPred, and Sitetack on AUC-ROC and AUC-PR for selected PTMs. Overall, AstraPTM matches or outperforms other methods in most common modifications (e.g., phosphorylation, glycosylation).

**Table 5:** PTM-specific **AUC-ROC** comparison for AstraPTM vs. MusiteDeep, ModPred, and Sitetack.

| PTM | AstraPTM | MusiteDeep | ModPred | Sitetack |
| --- | --- | --- | --- | --- |
| Phosphoserine/threonine | 97.95% | 89.6% | 75.3% | 92.80% |
| Phosphotyrosine | 97.95% | 95.8% | 69.5% | 90.10% |
| N-linked glycosylation | 99.68% | 99.3% | 77.4% | 98.90% |
| O-linked glycosylation | 94.40% | 94.3% | 78.3% | 96.00% |
| N6-acetyllysine | 97.59% | 97.8% | 70.2% | – |
| Methylarginine | 97.72% | 94.1% | 77.0% | 94.50% |
| Methyllysine | 97.72% | 95.1% | 67.0% | 96.80% |
| S-palmitoylation-cysteine | 99.46% | 96.1% | 82.4% | – |
| Pyrrolidone-carboxylic-acid | 99.99% | 97.9% | 86.0% | – |
| Ubiquitination | 94.27% | 80.4% | 58.4% | 92.30% |
| SUMOylation | 98.34% | 99.0% | 74.0% | 96.60% |

**Table 6:** PTM-specific **AUC-PR** comparison for AstraPTM vs. MusiteDeep, ModPred, and Sitetack.

| PTM | AstraPTM | MusiteDeep | ModPred | Sitetack |
| --- | --- | --- | --- | --- |
| Phosphoserine/threonine | 93.67% | 32.9% | 13.4% | 92.20% |
| Phosphotyrosine | 93.67% | 86.4% | 15.1% | 86.90% |
| N-linked glycosylation | 96.84% | 93.7% | 26.4% | 98.50% |
| O-linked glycosylation | 24.67% | 53.9% | 12.8% | 94.40% |
| N6-acetyllysine | 61.20% | 85.8% | 12.7% | – |
| Methylarginine | 46.50% | 84.4% | 13.0% | 91.00% |
| Methyllysine | 46.50% | 85.0% | 10.8% | 91.70% |
| S-palmitoylation-cysteine | 59.56% | 92.2% | 47.8% | – |
| Pyrrolidone-carboxylic-acid | 90.56% | 94.7% | 57.8% | – |
| Ubiquitination | 64.80% | 27.9% | 9.1% | 86.20% |
| SUMOylation | 26.42% | 88.1% | 21.3% | 94.50% |

### 7.2 Micro- and Macro-Averaged Performance

Table 7 compares micro-average (instance-weighted) and macro-average (class-weighted) performance across all PTMs for AstraPTM, MusiteDeep, and MIND-S.

**Table 7:**
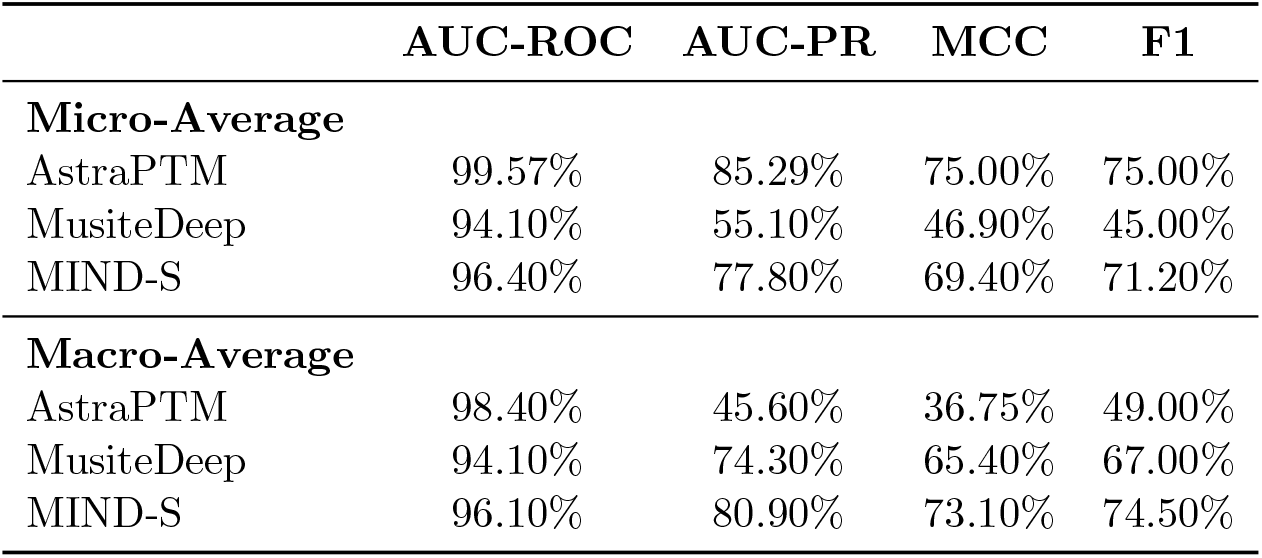
Micro- and macro-average performance (AUC-ROC, AUC-PR, MCC, and F1).

#### Micro-Average Superiority

AstraPTM leads in AUC-ROC (99.57%) and AUCPR (85.29%), confirming strong performance on frequent PTMs.

#### Macro-Average Considerations

Lower macro-average AUC-PR and MCC indicate room for improvement on rare modifications.

## 7.3 Comparison with MIND-S

### Insights from the MIND-S vs. AstraPTM Comparison

#### High AUC Across Most PTMs

AstraPTM achieves high AUC values on almost all modifications, indicating robust overall discrimination despite occasional drops in AUPR or MCC.

#### Trade-off in Precision and Recall

MIND-S sometimes secures higher AUPR (e.g., hydroxylysine (K)), suggesting it can excel when focusing on fewer false positives. AstraPTM, on the other hand, may capture more true positives but also more false positives, leading to variable AUPR in specific cases.

#### Variability in Rare PTMs

Both methods struggle on certain rare or highly imbalanced modifications (e.g., O-linked glycosylation with a low AUPR in AstraPTM), underscoring the difficulty of modeling less-characterized PTMs.

### 7.4 Comparison with PTMGPT2

We also compare AstraPTM to PTMGPT2 in Table 9, showing that AstraPTM often leads on major modifications like phosphorylation and glycosylation, while PTMGPT2 can perform better on certain low-frequency modifications.

### 7.5 Concluding Remarks

#### High Accuracy for Major PTMs

AstraPTM consistently achieves top-tier metrics for phosphorylation, glycosylation, and other high-frequency modifications.

#### Single-Model Multi-Label Efficiency

Users can streamline their pipelines without deploying separate binary models per PTM.

#### Room for Improvement on Rare PTMs

Like other methods, AstraPTM’s performance dips on heavily imbalanced or less-characterized modifications (e.g., lactylation).

#### Scalability Advantage

The transformer-based design accommodates full-length sequences, a key benefit for structural biologists studying large, multi-domain proteins.

While AstraPTM’s macro-average metrics indicate the need for improvement in certain rare PTMs, its strong performance on major modifications confirms its value for proteome-wide annotation tasks. The additional comparison against MIND-S (Table 8) further illustrates AstraPTM’s ability to maintain high AUC across diverse modifications, albeit with some trade-offs in precision-recall balance.

**Table 8:** PTM-specific performance comparison (AUPR, AUC, F1, and MCC) for MIND-S vs. AstraPTM.

|  | AUPR |  | AUC |  | F1 |  | MCC |  |
| --- | --- | --- | --- | --- | --- | --- | --- | --- |
|  | MIND-S | AstraPTM | MIND-S | AstraPTM | MIND-S | AstraPTM | MIND-S | AstraPTM |
| Hydroxylysine (K) | 89.20% | 61.25% | 97.60% | 99.40% | 81.10% | 64.00% | 79.60% | 62.52% |
| Hydroxyproline (P) | 55.20% | 61.25% | 89.10% | 99.40% | 47.30% | 64.00% | 40.00% | 62.52% |
| Methylation (K) | 87.20% | 46.50% | 99.30% | 97.72% | 69.60% | 47.00% | 72.50% | 41.83% |
| Methylation (R) | 63.20% | 46.50% | 95.10% | 97.72% | 65.50% | 47.00% | 65.00% | 41.83% |
| N6-acetyllysine (K) | 82.40% | 61.20% | 95.70% | 97.59% | 76.00% | 62.00% | 74.50% | 48.78% |
| S-Palmitoylation (C) | 94.70% | 59.56% | 99.30% | 99.46% | 89.50% | 53.00% | 87.90% | 53.31% |
| Phosphorylation (ST) | 64.20% | 93.67% | 94.60% | 97.95% | 59.80% | 90.00% | 58.10% | 85.46% |
| Phosphorylation (Y) | 73.90% | 93.67% | 92.40% | 97.95% | 67.30% | 90.00% | 64.50% | 85.46% |
| Pyrrolidone-carboxylic-acid (Q) | 90.40% | 90.56% | 97.30% | 99.99% | 80.50% | 84.00% | 76.90% | 81.35% |
| SUMOylation (K) | 96.50% | 26.42% | 99.60% | 98.34% | 87.10% | 31.00% | 86.90% | 25.87% |
| Ubiquitin (K) | 65.20% | 64.80% | 89.90% | 94.27% | 59.10% | 60.00% | 60.20% | 54.30% |
| N-linked glycosylation (N) | 92.80% | 99.68% | 99.20% | 96.84% | 90.30% | 95.00% | 89.00% | 94.75% |
| O-linked glycosylation (ST) | 96.20% | 24.67% | 99.80% | 94.40% | 95.40% | 34.00% | 95.40% | 17.94% |

**Table 9:** PTMGPT2 vs. AstraPTM on F1, MCC, Precision, and Recall for diverse PTMs.

| PTM | F1 |  | MCC |  | Precision |  | Recall |  |
| --- | --- | --- | --- | --- | --- | --- | --- | --- |
|  | PTMGPT2 | AstraPTM | PTMGPT2 | AstraPTM | PTMGPT2 | AstraPTM | PTMGPT2 | AstraPTM |
| Methylation (R) | 81.32 | 47.0% | 80.51 | 41.83% | 95.14 | 54.0% | 71.01 | 41.0% |
| Succinylation (K) | 60.74 | 38.0% | 51.88 | 9.20% | 79.15 | 34.0% | 49.27 | 44.0% |
| Sumoylation (K) | 69.21 | 31.0% | 66.93 | 25.87% | 75.87 | 29.0% | 63.63 | 32.0% |
| N-linked Glycosylation (N) | 74.21 | 95.0% | 70.92 | 94.75% | 64.35 | 97.0% | 87.65 | 93.0% |
| Methyl Arginine (R) | 81.32 | 47.0% | 80.51 | 41.83% | 95.14 | 54.0% | 71.01 | 41.0% |
| Methyl Lysine (K) | 36.35 | 47.0% | 41.07 | 41.83% | 88.68 | 54.0% | 22.86 | 41.0% |
| Lysine Acetylation (K) | 40.51 | 62.0% | 22.09 | 48.78% | 96.06 | 56.0% | 25.67 | 68.0% |
| Ubiquitination (K) | 35.74 | 60.0% | 31.25 | 54.30% | 80.46 | 52.0% | 22.97 | 70.0% |
| O-linked Glycosylation (S,T) | 61.80 | 34.0% | 58.97 | 17.94% | 58.79 | 32.0% | 65.14 | 36.0% |
| S-Nitrosylation (C) | 80.49 | 44.0% | 73.61 | 16.89% | 84.30 | 37.0% | 77.01 | 56.0% |
| Malonylation (K) | 78.14 | 24.0% | 73.03 | 0.00% | 82.95 | 19.0% | 73.85 | 35.0% |
| Phosphorylation (S,T) | 15.35 | 90.0% | 16.37 | 85.46% | 9.14 | 85.0% | 46.97 | 94.0% |
| Phosphorylation (Y) | 46.98 | 90.0% | 48.83 | 85.46% | 30.95 | 85.0% | 97.51 | 94.0% |
| Glutathionylation (C) | 82.37 | 44.0% | 78.28 | 10.13% | 90.07 | 37.0% | 75.89 | 53.0% |
| Glutarylation (K) | 73.78 | 19.0% | 69.47 | 0.00% | 71.02 | 26.0% | 76.76 | 15.0% |
| Amidation (V) | 76.59 | 93.0% | 80.78 | 93.10% | 66.67 | 92.0% | 86.76 | 95.0% |
| S-Palmitoylation (C) | 48.34 | 53.0% | 41.37 | 53.31% | 42.73 | 58.0% | 55.64 | 49.0% |
| Proline Hydroxylation (P) | 92.30 | 64.0% | 89.89 | 62.52% | 96.18 | 68.0% | 88.73 | 61.0% |
| Lysine Hydroxylation (K) | 68.18 | 64.0% | 66.45 | 62.52% | 88.23 | 68.0% | 55.56 | 61.0% |

## 8 Learnings

We introduced over 80 changes to architecture and hyperparameters, transitioning from a simple MLP to a BiLSTM and finally to a Transformer. This progression helped capture increasingly complex sequence patterns and improved overall metrics (Precision, Recall, F1, AUC–ROC).

### Model-specific Optimizations

#### MLPs

Lowering the learning rate (from 10^*−*3^ to 10^*−*4^) and tuning negative sampling rates pushed F1 from around 70% up to 85–88%. However, MLPs struggled with long-range context, leading us to explore recurrent models.

#### BiLSTMs

Integrating sequence context (hidden sizes 256–512, 1–2 layers) further improved recall beyond 90%, with F1 typically reaching 88–90%. We employed focal loss (*α, γ*) and dropout (0.2–0.4) to maintain precision.

#### Transformers

Ultimately, multi-head self-attention (4–8 heads) enabled learning of longer-range dependencies. With proper tuning of dropout (0.1–0.2), positional encodings, and thresholding, F1 climbed near 90%, with AUC–ROC at 95–96%.

### Context-Awareness Improvements

- BiLSTMs and Transformers surpassed MLPs primarily due to better modeling of sequence context; the former uses recurrent hidden states, while the latter uses self-attention to connect distant residues.
- For Transformers, refining positional encodings was crucial to improve alignment in the sequence, boosting both precision and recall significantly.

### Hyperparameter Optimizations

#### Threshold Tuning

Small changes in the decision threshold (48–50%) often led to large swings in Precision and Recall, highlighting its importance for imbalanced data.

#### Focal Loss and Sampling

Varying negative sampling rates (1.0–3.0) and carefully setting focal loss (*α, γ*) helped reduce false negatives while avoiding too many false positives.

### Model Dimensions and Data Growth

- Increasing hidden dimensions (256 to 512, or higher for Transformers) generally elevated F1 by 1–2%, though it required more GPU memory and longer training time.
- Expanding our dataset from 40K to 94K examples consistently improved all architectures, provided the sequences were diverse and representative of real-world PTM patterns.

### Key Takeaways

- A well-tuned **Transformer** with focal loss, moderate dropout, and balanced negative sampling produced the best F1 (near 90%) and highest AUC–ROC (95–96%).
- Careful hyperparameter scheduling (e.g., warm-up, OneCycleLR) and threshold calibration were critical to maintaining stable convergence and robust final metrics.
- Addressing class imbalance with negative sampling and focal loss had a stronger impact than any single architectural change alone.

Overall, the transition from MLPs to sequence-aware models (BiLSTMs, Transformers), combined with thorough hyperparameter optimization, was instrumental in achieving state-of-the-art performance for this PTM prediction task.

## 9 Next Steps

Although AstraPTM demonstrates strong potential for identifying PTMs and suggesting likely modification sites, further work is required to (1) boost accuracy/performance for rare modifications and (2) improve usability for end users.

### Enhancing Model Accuracy and Performance

#### Integrate AlphaFold Structure Predictions

PTMs are heavily influenced by 3D conformation. Incorporating structural predictions could provide crucial spatial context.

#### Increase Training Data

*dbPTM* was used as the primary source. Supplementing with other databases or curated mass spectrometry datasets may help mitigate data imbalance for rarer PTMs.

#### User-Friendly Interface

A web-based or desktop application with interactive visualization can help researchers easily interpret PTM predictions across large protein structures.

